# Exploring the Potential of Large Language Models in Differential Abundance Analysis

**DOI:** 10.1101/2025.06.04.657820

**Authors:** Roberto Franco-Alba, Irina Goryanin, Igor Goryanin

## Abstract

The rapid development of Large Language Models (LLMs) has opened new possibilities in various fields, including microbiome research. This dissertation explores the application of LLMs for differential abundance analysis, a crucial method for understanding the relationship between microbial communities and health conditions. Through a series of experiments, we assessed the capabilities and limitations of LLMs in automating and enhancing the differential abundance analysis process.

Our findings reveal that while LLMs can effectively extract relevant information from scientific literature and assist in generating reports, they also face significant challenges, including data inaccuracies, hallucinations, and issues with reference generation. These challenges highlight the importance of integrating LLMs with human oversight to ensure scientific rigor. The study suggests that while LLMs can increase research efficiency, they are not yet reliable enough to replace human expertise in complex scientific tasks.

This research contributes to the ongoing dialogue on the role of AI in scientific research, emphasizing the need for further development of LLMs and the exploration of hybrid models that combine AI capabilities with human judgment. The dissertation concludes with recommendations for improving the accuracy and consistency of LLMs, and expanding their applications.

## Introduction

LLMs are a type of artificial intelligence trained on massive datasets of text and code, enabling them to generate human-quality text, translate languages, and write different kinds of creative content [32]. They are currently an area of very active research, with several industry applications that range from narrative generation and chatbot engineering to music and image creation [1, 3, 6, 21, 2, 34, 28]. In recent years, the integration of computational tools with biological research has markedly transformed the landscape of microbiology; from identification of genes and gene functions to regulatory elements within microbial genomes [29]. Microbiology underpins a vast swathe of scientific inquiry, from understanding human health and disease to bioremediation and industrial processes. A cornerstone of modern microbiology is the analysis of microbial communities, often through next-generation sequencing technologies [23]. Differential abundance analysis (DAA), a critical method in microbiomics, identifies species that are differentially abundant across various conditions or environments, providing insights that are pivotal for understanding microbial communities and their functions, and allowing researchers to identify microbes that are statistically different between two or more conditions [8, 37, 30, 20]. This has yielded a data deluge, with researchers grappling with the challenge of extracting meaningful information from data-intensive analyses such as DAA [8]. Also, translating this data into clear and concise scientific narratives remains a time-consuming bottleneck [41, 4]. This dissertation aims to explore the potential of LLMs as a novel tool to engage in scientific research by addressing this bottleneck.

In Background we describe the current state of the art in LLM capabili ties, and we summarize the most recent findings in the crossroads between LLMs and Scientific Research (including Biology Research), we also provide a brief description of the typical process to perform differential abundance analyses.

In Methodology we describe the process by which our approach, data sources, and precise objectives were set, and we describe the script’s interface with the user as well as the architecture of our report-generation and our output-evaluation scripts. We also describe the methods used to reach our conclusions.

In Evaluation we describe the characteristics of the output obtained at each one of the steps of the report-generating process, as well as the evaluation of output against their expected forms.

In Discussion we describe the implications of the evaluation findings, we elaborate on the study’s limitations, and provide suggestions for future research.

Finally, in Conclusions we summarize the main findings of the study and propose a path forward towards improving the process for an automated differential abundance analysis.

## Background

### 2.1 What are Large Language Models?

“A language model is a probability distribution over sequences of words” [3]. In turn, LLMs are at the core of solutions including “machine translation, speech recognition, text generation, sentiment analysis[, etc.]” [3]. Generative LLMs are LLMs that are able to automatically complete text that is likely to follow a prompt provided by a user. This would be done on the basis of the patterns learned from reading vast amounts of literature [3]. Contemporary generative models (such as BERT or GPT-4) have achieved strong performance in a number of tasks [28], and are capable of creating different types of output for immediate user consumption. From its theoretical starts in the second half of the 20th century, a number of algorithms have been created to address specific issues related to the challenge of working with text as data: from vector space models (SVM) in the late 1960s for information retrieval, to the distributed representation of words to encode grammatical and semantic information, to word embeddings to capture the diverse contexts for each word [39]. Generative Pre-Trained (GPT) models were first proposed by Radford et al. (2018) [31], based on a recent development by Vawanit et al. (2017) [35], by which a model would be able to encode and decode text by paying attention to different parts of the input sequence at the same time. This made the algorithm ideal for tasks such as text generation and language translation. On top of this, a number of incremental releases of the GPT model have provided it with ever-increasing generative capacities [28]. GPT-4 has purportedly exhibited human-level performance in several standardised tests [28]. However, its ability to generate scientific knowledge based on formal descriptions of microbial communities, to the extent of our knowledge, had not been tested. Generative models have been documented to generate text for a wide variety of purposes, including those of a very technical nature: technical documentation of construction updates for railway overhead contact systems [36]; OpenAI’s recent release of a ”Data Analyst” GPT ^1^, which provides powerful data description capabilities; and Microsoft Power BI’s ”describe the data” feature [24], which generates short sentences that describe a given dataset [25]. Nonetheless, it is unclear whether the models’ current capabilities allow for a streamlined generation of knowledge from publicly available data.

### 2.2 Application of LLMs in Scientific Research

LLM’s potential in the scientific community has been of particular interest, given their ability to process and synthesize large amounts of complex scientific literature and data quickly. This ability positions LLMs as powerful tools for accelerating scientific discovery, assisting with data analysis, hypothesis generation, and even direct interaction with experimental setups [5, 2, 21]. The application of LLMs in scientific research has been explored across various domains such as drug discovery, biology, computational chemistry, materials design, and solving partial differential equations [2]. In drug discovery, LLMs like GPT-4 have shown promise in tasks such as drug-target binding prediction, molecular property prediction, retrosynthesis, and novel molecule generation [6]. These capabilities suggest that LLMs can significantly reduce the time and cost associated with the early stages of drug development by predicting potential drug candidates’ efficacy and side effects before they are synthesized [2]. In biology and materials design, LLMs have been utilized to understand complex biological sequences and assist in the design of new materials by predicting properties and synthesizing routes for novel materials. Such applications underscore the models’ ability to enhance productivity and innovation in these fields [2]. Also, systems of LLMs have been found to be capable of conducting sophisticated searches, controlling laboratory hardware, and performing experiments autonomously [6]. That is, LLMs were capable of planning, coordinating, and summarizing research. Finally, LLMs have been found to generate useful feedback on scientific papers that are often evaluated better than those produced by humans [18]. It is thus suggested that LLMs can be especially helpful for researchers in the early stages of manuscript preparation, and for those who might lack access to detailed or prompt peer reviews [18]. However, LLMs have not often been described as unable to produce original scientific contributions, which highlights a significant limitation in their current use in academic settings [21].

### 2.3 LLMs in Biology Research

In recent years, the integration of computational tools with biological research has markedly transformed the landscape of microbiology; from identification of genes and gene functions to regulatory elements within microbial genomes [29]. Moreover, LLMs have been recently used for the following use cases:

- for predicting antibiotic resistance and associating microbiome features with complex diseases [4],
- for “summarizing laboratory reports (eg, key findings, abnormal findings, notes on uncommon pathogens, advice on antibiotic selection, and next steps in testing)” [11],
- for generating “protein sequences with predictable functions across large protein families” [22].
- for analyzing “genomic data to predict functional categories learn the evolutionary landscape of 1.5 million severe acute respiratory syndrome coronavirus 2 genomes and accurately and rapidly identify variants of concern” [26].
- for enhancing the annotation of viral proteins from prokaryotes in metagenomic data, classifying viral proteins, and predicting functions of viral proteins beyond what is possible with traditional methods [13].
- for “analyzing and interpreting microbial genomic data” [19].

As the above examples illustrate, LLMs have emerged as powerful assets, promising a shift in how data is processed and interpreted in scientific research. The utility of LLMs extends beyond simple text generation; they are increasingly being harnessed to distill complex datasets into comprehensible, insightful academic literature.

### 2.4 Differential Abundance Analysis

Differential abundance (DA) analysis involves comparing the microbial communities between different environments or conditions to identify taxa that are significantly different in abundance [20].

It has been found that” a stable gut microbiota is essential for normal gut physiology and overall health” [33] and has a direct impact not only in the brain-gut axis but also on the immune and other systems [12] and may hold the key to unlocking better understanding and treatment of a number of health conditions. [10]

Therefore, understanding the variations in abundance of specific bacteria species across groups of people with different health characteristics may be at the core of important discoveries soon. Nonetheless, it has been found that different DAA methods can produce different results, and often the amount and variability of data make it complex to draw conclusions [27]. Therefore, we believe that the task of procuring data, processing it, and comparing different findings from similar studies to reach more robust conclusions is central to understanding the relationship between microbiota and specific health conditions. We also believe that LLMs may possess the technical characteristics that are needed to explore large amounts of data and draft meaningful conclusions from it. This complements the need to reduce the time and effort needed to generate such results.

A common path to perform a differential abundance analysis would include the following:

- **Data Collection**: Collecting stool samples from healthy and diseased subjects.
- **DNA Sequencing**: Performing 16S rRNA gene sequencing on the collected samples.
- **Quality Control**: Removing primers, demultiplex, and filter sequences.
- **OTU Construction**: Clusterising sequences into OTUs using UPARSE.
- **Normalization**: Applying Cumulative Sum Scaling (CSS) to normalize the feature table.
- **DA Testing**: Using ANCOM-BC to identify differentially abundant taxa.
- **Result Interpretation**: Generating p-values, confidence intervals, and visualizations to report findings. [20]

Due to the variability in data formats, as well as the heterogeneity in methods used (and the complexity and compute time required to pre-process data close to the DNA sequencing step), we aim to gather data from published sources that are at least at the **DA testing** stage.

## Methodology

### 3.1 Program and Methodology

Having in mind the general objective of achieving an automated differential abundance analysis, we set specific objectives that evolved in time while we learned more about the data available in usable data sources, about the fast-evolving capabilities of commercial engines such as Gemini ^1^ and ChatGPT ^2^, and about the initial results observed during the exploratory phase.

A first plan involved extracting raw metagenomic data from an open repository such as MG-RAST ^3^ or MGnify ^4^ and producing a written output. However, data available in both repositories required important pre-processing (to bring it from a raw or DNA sequencing format to a DA testing stage) and often did not contain enough metadata to link individual patient observations to control and condition groups. Missing metadata was often found in separate repositories (such as Supplementary Information to published research papers in academic journals), but its format ^5^ and completeness varied considerably from case to case, which made automated data pre-processing extremely complex. Also, manually gathering relevant data for development purposes required more subject-matter expert involvement than was available and was also considered contrary to the original idea of producing a script that would be capable of automated analysis and report writing with minimal end-user involvement.

For the above reasons, it was decided that data would be sourced from research papers published in academic journals. Data in research papers is at times included in the body of the document or as supplementary data (which could have various forms but would typically be pre-processed, normalised, and conforming to a DA testing standard -as opposed to a raw or DNA sequencing format or at a sample accession granularity-). That is, available data would be easier to consume and re-utilise without much additional pre-processing.

Once the source of the data was decided, we explored the possibility of triggering the entire extraction, analysis, and drafting mechanism using a single prompt to an LLM engine. That is, providing the LLM a single paragraph with a detailed list of instructions, and exploring the generated output. This approach offered the benefit of simplicity and the lack of any additional software requirements. However, generated reports were full of inconsistencies and hallucinations ^6^, and also, a single-prompt approach didn’t allow us to inspect where in the internal ”thought” process of the black box the process started to get lost.

The next natural step was to type each prompt separately and sequentially to the LLM chat and evaluate the output at each stage. Starting with an instruction to get a list of relevant articles, then progressing to data ingestion, and so on. This option also allowed us to dialogue with the LLM and request corrections to wrong or inaccurate outputs. This approach was followed by Caspi and Karp (2024) [9].

However, it was soon found that both of our commercial LLM engines (Gemini and chatGPT) were not a good source for our initial list of articles: both engines consistently returned an insufficient number of entries and very often returned references that were partially or completely hallucinated. Also, and in line with the findings of Caspi and Karp (2024) [9], no subsequent dialogue was productive in making the LLMs return a complete list of correct entries.

It was deemed imperative that the initial data exploration and ingestion steps returned accurate information, to avoid very early error propagation to all subsequent steps of the process which would then magnify the scale of hallucination. For this reason, we abandoned the chat dialogue in favour of a Python script.

Our Python prototype would use web-scraping capabilities to extract lists of research articles from the sources themselves. The script would use a rules-based approach to filter a preliminary list of articles and then feed them sequentially to LLMs for additional transformations.

Finally, the full process was set to include the following steps:

1. User requirements gathering
2. Data exploration (web-scraping)
3. Data ingestion
4. Table consolidation
5. Report Generation

#### 3.1.1 User Requirements Gathering

As per the initial design requirements, an end-user would initialise the program by providing the following pieces of information:

- **The name of a health condition**: a human health condition (such as *hypertension*) that would serve as a keyword to search for relevant research papers in research journals. The web-scraping function would initially search for articles about the provided health condition that also mentioned ’microbiome’ or a related term.
- **An initial number of research papers**: the ideal number of research papers to use as a basis for the analysis and subsequent report. These would have to contain all the necessary data elements. At a later stage, the script extracts bacterial data from these research papers. In the final version, the script was set to draft a report even if it finds fewer papers than requested (as long as some data is retrieved). The user is notified about the number of relevant papers found. Also, the script stops downloading papers upon reaching the required number.
- **A URL to an ’inoculum’ paper**: A research paper with material from which to start the analysis and drafting process.
- **A URL to a style guide**: Typically the style guide or the publishing requirements page of a scientific journal. This page would provide a guideline of the expected standard for all formal elements that need to be included in the final report.

#### 3.1.2 Data Exploration

During the Data Exploration phase, the script was planned to query various scientific journals. From each journal, it would extract a list of relevant articles which would be reviewed at a later stage. After discarding MG-rast and MGnify during an earlier exploratory stage (for having data in raw or DNA sequencing format, thus requiring considerable pre-processing), all of the below journals were evaluated as data sources for the project:

**Table 3.1:** Evaluation of possible data sources. Ease of access: 1-Very simple; 2 - Simple; 3 - Difficult; 4 - Very difficult.

| Source | Microbiome data | Metadata available | Full articles | Robot-friendly |
| --- | --- | --- | --- | --- |
| **bioRxiv** | 4 | 4 | 1 | 1 |
| **Nature** | 2 | 2 | 1 | 1 |
| **Springer** | 2 | 2 | 1 | 1 |
| **Science Direct** | 2 | 2 | 1 | 4 |
| **Wiley** | 2 | 2 | 1 | 4 |
| **MDPI** | 2 | 2 | 1 | 1 |
| **Microbiology Research** | 2 | 2 | 1 | 1 |
| **Biomedcentral** | 2 | 2 | 1 | 1 |
| **Taylor & Francis** | 2 | 2 | 1 | 4 |
| **American Society for Microbiology** | 2 | 2 | 1 | 4 |
| **Cambridge** | 2 | 2 | 1 | 1 |
| **PLOS** | 2 | 2 | 1 | 4 |

From the above, the following sources were ultimately chosen: Nature, Microbiology Research, MDPI, Biomed Central, and Cambridge.

The most important requirement to consider was the ability to robotically query journals, according to the search parameters specified by the user (i.e. a health condition, such as Hypertension). This is because some journals would identify robotic users (as opposed to human ones) and decline to send queried information.

Also, some of the above data sources were favoured because microbiome data was available in the right format and without the need to perform additional or complex steps to download it (biorXiv ^7^, for example, pointed users to external sources to fetch data; often, users would need to click their way in these external sources to find and download the data).

Overall, for the 5 chosen sources, data required little to no pre-processing to get an output that would tell a story from an experimental point of view (that is, grouping results by illness or health group, or treatment and control group). Nonetheless, even in our final set of article sources, supplementary materials were found to vary widely in both quality and format in each case, and this meant we had to consider an additional step in our script to explore each article and its data to determine their eligibility.

For this purpose, we used LLM calls to review the contents of each research paper and make sure it contained all the required elements:

- Supplementary Information
- Information about bacteria species
- Information about gene names
- Experimental information on the health condition (such as hypertension) and control groups.

An additional requirement to contain information about phylogeny, although highly desirable, was removed from the process because most of the articles did not contain it, leaving us with no data to work.

The process would sequentially prompt GPT-3.5 to review each article and evaluate whether it contained each one of the above elements. GPT-3.5 was used because newer models very often declined to review URLs that we provided. The Python script would then flag articles that contained all required elements and stop evaluating more once the number of relevant articles reached the number parameter set by the end user at the beginning, or when the full list of articles retrieved from journals was exhausted.

The list of relevant articles was then sent to the next stage of the script for data ingestion.

#### 3.1.3 Data Ingestion

Information extraction was done in the form of tabular data extraction. Relevant data can be typically found in publications under different formats and places: in tables within the body of the report, or in external files contained in the Supplementary Information, etc.

For this reason, it was decided that LLM involvement would be instrumental in deciding where data was found, and extracting it.

Our data ingestion function receives a list of relevant article URLs and returns a data table for each entry in the list. The function does this by sequentially sending each article to an LLM, together with a prompt. The prompt requests the model to scan the article found under the provided URL (and any available supplementary information) for microbiome data and generate a table following a specified format.

The prompt requests the LLM to appropriately mark with NA any information that is not found in the document (to prevent as much as possible the addition of hallucinated elements from the model’s pre-trained information into the tables).

Due to the length of some of the articles, and to the large number of files that are sometimes found in the supplementary information, and knowing that LLMs have a limited context length ^8^, we thought of aiding the models to find relevant data by adding a semantic search functionality to the script.

This ideal solution would have included an additional step to download all relevant data for each research paper and, using a microbiology-trained embeddings model, split documents into semantically homogenous chunks, create a vector search index, use the index to search for relevant data within each document and then feed the LLMs only the relevant chunks in each case. However, budget considerations (needed to set up the appropriate architecture) meant that this option was not viable.

Our solution thus leverages the commercial models’ internal tools to extract data and meaningful information from any submitted text. Although their internal architecture and specifics remain largely a mystery to the final user, the current industry standards and achieved model accuracy suggest that commercial engines should utilize some type of chunking and indexing solution of uploaded files and retrieved internet pages to feed their models the most appropriate content. However, we believe that these internal indexing tools are multi-purpose and thus unlikely to be tailored to microbiology jargon. Hence the possibility of improving accuracy by extracting data using a model fine-tuned for microbiology language.

Our data extraction mechanisms consist of a single-shot prompt that was fine-tuned to minimize output variability and facilitate ulterior algorithmic processing of the received tables.

All generated tables (which contained bacterial information from research papers) were then fed to an LLM together with a prompt to have them combined into a single table.

This step was performed using an LLM because of the observed variability in the individual tables created from each research paper. That is, our single-shot prompt reduced output variability but still produced tables with slight style and format differences every time. These were easier to consolidate into a single table using an LLM than a rules-based algorithmic approach.

#### 3.1.4 Table Consolidation

A simple LLM prompt, along with all individual tables, was used to get our final consolidated table.

As discussed above, a simple concatenation did not suffice due to the slight variability observed in individual tables (even as we provided examples to the LLM to reduce format variability). Observed variations included padding differences, row-separation lines, and other creative usage of column separators.

Upon consolidation, the final table would typically have a uniform format that looked consistent and readable.

#### 3.1.5 Report Drafting

Once all data is consolidated into a single table (which contains traceability information on each row, to follow each finding to its original published source), we used an additional prompt to feed the table to the model and request a written report with specific characteristics.

The required characteristics included:

- It should describe any novel findings based on the ”inoculum” paper and the provided data.
- It should follow an academic format with formal elements such as an abstract, introduction, background, findings, discussion, conclusions, and bibliography.
- It should include citations (recent bibliography preferred).
- It should include tables with any relevant data.
- It would follow stylistic guides from a specific scientific journal.

The final report was drafted following two different processes. The first was a single-prompt, single-call generative process, and the second was created in parts and stitched together. The idea behind this is the following: a single call would be able to return a clear and coherent report within the maximum token count allowed by the model; the multi-call process would allow the LLMs to elaborate further and in more depth in each section, but would risk having each chunk develop in different directions.

*Appendix A.1* has a list of all prompts used for creating the two versions of the final report.

#### 3.1.6 Evaluation

The output of each specific research objective is evaluated separately, and a script has been created for this. The next chapter describes in detail how the evaluation was planned and executed.

## Evaluation

The output of each step is evaluated separately. For this purpose, an evaluation script was created and both scripts were run for five different health conditions. However, not all output could be evaluated in an exclusively automatic fashion. Although a higher number of runs against a wider set of health conditions would have benefited the study’s objectives, the number of runs was constrained by the amount of output and throughput elements that could be manually reviewed.

The evaluation methods and outputs will be presented within the structure of the overarching generative script. In accordance with the above, the entire evaluation process consisted of the following steps:

- User provides run parameters.
- Automated Data Exploration.
- Automated Data Ingestion.
- Automated Tables Consolidation.
- Automated Report Generation.

### 4.0.1 Initial run parameters

An automated working pipeline was designed to take any health condition as input from the final user and produce an output (provided there would be data that would be relevant to such condition).

Nonetheless, for practical purposes, we circumscribed our initial study to five main conditions for which enough data was expected to be available. Below is a table containing the parameters used to run the script in each run. For each one of the below, final reports were generated, as well as all expected throughput files for evaluation.

**Table 4.1:** Evaluation run parameters.

| Run | Condition | # of articles | "Inoculum" paper | Style guide |
| --- | --- | --- | --- | --- |
| 1 | Hypertension | 15 | See [17] | Medinformatics [15] |
| 2 | Rheumatoid Arthritis | 15 | See [38] | Medinformatics [15] |
| 3 | Chronic Obstructive Pulmonary Disease | 15 | See [7] | Medinformatics [15] |
| 4 | Crohn's disease | 15 | See [16] | Medinformatics [15] |
| 5 | Irritable Bowel Syndrome | 15 | See [14] | Medinformatics [15] |

Parkinson’s disease was originally also used as input, but later removed from the study, as the script found no relevant articles with workable data.

The choice of health conditions consisted of an initial triad provided by subject matter experts, further complemented by an additional two for which sufficient data was available.

The ”inoculum” or initial papers for each condition were hand-chosen for their completeness and relevance and were used as initial examples to trigger text generation and provide initial bacterial data for the review.

The style guide was the submissions guidelines page of the Medinformatics journal. Its purpose was to provide the model with information about expected formal elements and style for the final report.

Finally, the number of articles was set to 15 and not higher due to the need to manually evaluate if models were extracting all available data from articles in the desired way.

For reproducibility purposes, all the above parameters and throughput were saved as Python pickle files and are available for download together with the script repository ^1^.

### 4.1 Data Exploration

The data exploration phase aimed to explore available data for completeness and choose the most appropriate datasets for eventual ingestion. To do this, the script queries five research journals for entries relevant to microbiome data and the health condition provided by the user. Later, each article from the initial list is inspected by an LLM to check whether it contains all required formal elements. All relevant articles are passed to the data ingestion phase to retrieve bacterial data from them. The evaluation task at this stage had the objective of measuring the following:

- The amount of available data and the best sources identified for our task.
- The distribution of required formal elements across all inspected articles.

#### 4.1.1 Candidate Articles

Table 4.2 shows the number of articles that were extracted for exploration from each research journal (for the five health conditions together), along with the number of articles that passed the evaluation filters and were used for data extraction.

**Table 4.2:**
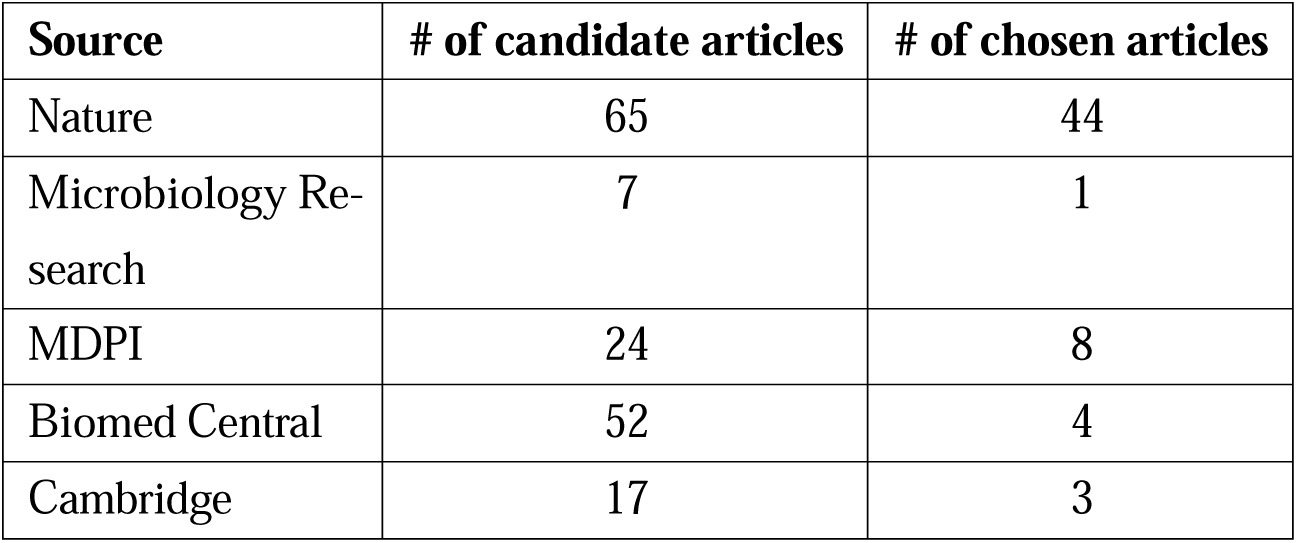
Sources of articles for ingestion.

| Source | # of candidate articles | # of chosen articles |
| --- | --- | --- |
| Nature | 65 | 44 |
| Microbiology Research | 7 | 1 |
| MDPI | 24 | 8 |
| Biomed Central | 52 | 4 |
| Cambridge | 17 | 3 |

The proportions of chosen versus candidate articles are not to be given excessive importance because the script would stop evaluating candidate articles once it found good articles to match the expected article count (which we set at 15 in every case).

Also, the order of appearance in the table 4.2 coincides with the order in which the script queries and evaluates the articles from each journal, which is the main reason why most chosen articles come from Nature. Nonetheless, it is worth noting that not all journals contributed equally to the list of article candidates.

Table 4.2 thus provides a reference of the best article sources relevant to microbiome research and human health conditions for any future research.

#### 4.1.2 Elements Found

The process of choosing articles from the candidate list implies having the LLMs read each document and determine if all the required elements were present. A manual review of the shortlisted articles suggests that over 90% of all chosen articles was topic-relevant. However, Table 4.3 shows the proportion of reviewed articles that were identified by models to contain each one of the required elements.

**Table 4.3:** Element prevalence in articles.

| **Element** | **% of chosen articles that contained it** |
| --- | --- |
| Bacterial species names | 73% |
| Gene names | 67% |
| Supplementary Information | 77% |
| Experimental Information | 78% |

Due to the substantial number of articles processed at each run, both those ultimately included or excluded from the pipeline, a manual review of the initial article evaluation phase’s accuracy was beyond the study’s scope. Nevertheless, we can compare the element-existence proportions in the table above to the actual elements retrieved. While extracted elements are examined in greater detail in the next subsection, an initial comparison reveals a significant discrepancy between the tables. The table above suggests that 78% of reviewed articles included experimental information on participants’ health status (ill vs. healthy, or treatment vs. control groups). Moreover, 100% of articles chosen for data extraction should contain the above information, yet in most cases, such data fields in our extracted tables were empty. A similar pattern is observed for gene function data.

In sum, the high proportions indicated in the table above do not align with the proportions observed in extracted data, pointing to the fact that LLMs failed to either properly flag the existence of required elements or to properly extract all available data from chosen articles.

### 4.2 Data Ingestion

An LLM inspected all short-listed articles to extract from each one of them a table with relevant data (including bacterial species, genes, etc.)

The evaluation task at this stage had the objective of measuring the following:

- The LLMs’ ability to extract all **relevant data** from short-listed articles.
- The **usefulness** and data richness of each **short-listed article**.
- The **quality of data extracted** from short-listed articles.

#### 4.2.1 Relevance of Extracted Data

Table 4.4 shows the proportion of bacterial species that appeared in the summary tables of short-listed articles and were also present elsewhere in the article or its supplementary data. It also shows the complement: the proportion of bacterial species that were not found anywhere in the articles or their supplementary data.

**Table 4.4:** Element prevalence in articles.

| Condition | % of bacteria species |  |
| --- | --- | --- |
|  | Rightly attributed | Wrongly attributed |
| Rheumatoid Arthritis | 10 (15%) | 57 (85%) |
| Hypertension | 12 (24%) | 38 (76%) |
| Crohn's disease | 25 (83%) | 5 (17%) |
| COPD | 47 (80%) | 12 (20%) |
| Irritable Bowel Syndrome | 47 (80%) | 12 (20%) |

The analysis of the bacterial summary tables extracted from individual articles allows us to confirm that hallucination was commonplace, although its prevalence varied significantly from one health condition to another.

The above suggests that the LLM we queried for our study (GPT-3.5), although capable of consulting sources from the internet, was not a reliable source for summary tables.

A subsequent test using GPT-4o’s chat capability (GPT-4o’ API often refused to consult sources on the internet, making it difficult to query it programmatically) showed GPT-4o is much more accurate in extracting bacterial data from articles, and in generating summary tables.

#### 4.2.2 Usefulness of Short-Listed Articles

A manual review of the short-listed articles for the presence or absence of expected elements suggests that several lacked one or more of the required elements. This points to the fact that, in at least 20% of the cases, GPT-3.5 falsely claimed that an article contained an absent (and required) element.

Further manual review of the same articles using GPT-4o’s chat showed an improvement in accuracy. Importantly, GPT-4o’s chat (as opposed to the same model queried through its API) did not decline a request to review the contents of a page when provided with a URL. GPT-4o still was unable to provide a good initial list of journal articles to kick-start our process.

#### 4.2.3 Quality of Extracted Data

An additional (manual) review of the tables and the original sources showed that the summary tables often omitted bacterial species that were central to the original source (in some cases even being mentioned in the article’s title); on the contrary, sometimes bacteria in the summary table was only mentioned in a bibliographic reference of the paper in question (and not in the body of the work, or the supplementary information); in other cases, bacterial species could be found in the body of the article but none of the additional elements present in the table (gene functions, biochemical pathways, etc.) were found to be present anywhere in the article; finally, bacteria that did appear in the original texts or supplementary tables, were not necessarily mentioned to present differential abundance or to be otherwise relevant in terms of conclusions.

For all of the above reasons, we believe that the quality of the data extracted by the LLMs in all five runs was unreliable and not a true representation of the information contained in the source articles. The best data extraction results were obtained by querying GPT-4o directly in the chat (yet, API calls to the same model would return a denial to review contents from the internet, thus limiting our possibility to integrate GTP-4o with our automated pipeline).

### 4.3 Table Consolidation

All tabular data extracted from each short-listed article was then consolidated into a single table. The evaluation task at this stage had the objective of measuring the following:

- **Completeness**: If all data captured in individual tables was correctly consolidated into the final table.
- **Accuracy**: If the consolidated table contained entries that did not appear in the initial tables.
- **Quality**: The overall quality of the data tables.

#### 4.3.1 Completeness

From the bacteria species present in individual tables, only 43% were included in the consolidated table. The percentage of bacteria species that made it from the individual table into the consolidated one varied in each study (from 31% for Rheumatoid Arthritis and COPD, to 60% for Crohn’s disease).

Looking at the same data tabulated by bacteria species suggests that different bacteria were treated differently. While the number of observations does not allow for robust statistical conclusions, the data does seem to suggest that some bacteria (such as *Bacteroides fragilis* and *Escherichia coli*) were never omitted, while other bacteria species (such as *Lactobacillus* and *Streptococcus*) were always left out during consolidation. Unfortunately, increasing the number of runs to produce more data would have made it difficult to perform manual data quality checks and to review output texts manually.

#### 4.3.2 Accuracy

Across all runs with different health condition parameters, all bacteria species in the consolidated tables also appeared in the individual tables. When the same bacterial species appeared across different individual tables, it is unclear what process was followed to summarise or choose bacterial attributes if they differed across individual table rows.

#### 4.3.3 Quality

Although bacterial species found in the consolidated tables mostly appeared in the individual tables, the details related to each species were often modified. DOI information was frequently hallucinated in the consolidated table, even when the correct DOI was present in the individual tables. Additionally, it is unclear what caused certain bacterial species to be omitted from the consolidated list, as both species with complete additional information and those with mostly missing values were excluded similarly.

The consolidated table seems to be, at most, an artistic representation of how such a consolidated table would roughly look like.

### 4.4 Report Generation

The consolidated data table was fed to an LLM to produce a report in two different ways: a) using a single, one-shot prompt and b) drafting various elements separately and stitching them together (to increase the report size and depth). The first report-drafting approach explored the possibility of creating a coherent report with minimal input, the second approach explored the models’ capabilities to elaborate deeper in each section. The two report types were evaluated separately. The evaluation task at this stage had the objective of measuring the following:

- The quality of included formal elements.
- Whether any important elements were missing.
- The quality and relevance of the bibliographic references.

For this task, a second LLM instance was used to evaluate outputs as there was not enough availability of subject matter expertise to review the outputs manually. Based on conclusions from Liang *et al.* (2023) [18], we determined that LLMs should be able to provide useful and accurate feedback on the reports’ quality.

#### 4.4.1 Single Shot Reports

All articles were first evaluated by a second LLM instance (Gemini) and then reviewed by the author for correctness. The LLM was prompted to list both the strengths and weaknesses of the attached report. Additionally, all reports underwent an additional evaluation of their bibliography and references due to the frequent occurrence of bibliography-related hallucinations.

In almost all cases, articles were praised for their clarity, concise titles and summaries, and clear acknowledgment of limitations. Most bibliographic references were praised as relevant and appropriate. However, observed shortcomings included a lack of visual aids, issues with the quality and completeness of references and provided data, a limited number of data sources, incomplete DOI references, and, in some cases, small sample sizes. In a few cases, bibliographic references were irrelevant or completely made up.

The lack of visual aids was understandable, as the final report, by design, did not contain them. Early exploratory research showed that GPT-3.5 and Gemini could only create artistic representations of visuals that did not accurately represent the provided data. Instead, a more accurate approach was to request Python scripts to generate such visuals. However, a pipeline that included these elements in the final report was not planned due to time constraints and the complexity of programmatically debugging Python scripts output by LLMs, which sometimes fail to run.

Regarding scientific rigor, some comments from the evaluating model included: limited discussion of clinical implications, bibliography that did not support central ideas or claims, bibliography that was not exhaustive or lacked key articles, issues with the data (such as important elements missing), and in one case, part of the bibliography resembling a template with placeholder elements to be completed later. Often, the lack of in-depth discussion or additional information was cited. We believe the main reason for this is the models’ context window, which importantly constrained the length of the generated reports.

Given the various flaws in the data-gathering process, it is assumed that any reported findings would have weak scientific validity. In some cases, the evaluating LLM instances also suggested this, citing the lack of supporting bibliographic evidence for central ideas in the report.

In summary, the reports scored highly for the quality of their writing, which is expected as our models (GPT-3.5 and Gemini) were optimized for text drafting. However, scientific rigor suffered, as expected standards in reference generation and supporting ideas were not met. This highlights that the models used were not optimized for scientific writing.

#### 4.4.2 Stitched Reports

Stitched articles were created by prompting our models six times (once for each section of the text: Abstract, Introduction, Discussion [queried twice for further elaboration], Conclusions, and Bibliography). These components were then stitched together, and the model was prompted again for style and cohesion. This approach allowed for a longer, more in-depth report, as the final query did not need to include a long, consolidated data table, which would have consumed the model’s context window, thereby limiting the length and depth of the final output.

In almost all cases, reports were praised for their clarity, their well-organised structures, and for being comprehensive and detailed in the analysis. In some cases, the evaluating model praised that the text identified areas for future research and described the findings as ”intriguing”, with potential for therapeutic interventions. The presented bibliography was widely described as relevant and appropriate, covering a wide range of topics.

On the other hand, the most common shortcomings included: the lack of data or insufficient data (understood as it being not reported in the report, or the sample size being too small), the lack of detailed methodologies and statistical analysis (understood as how gene expression or other elements were studied across different bacterial species), the lack of visual aids (which was expected, as with the single-prompt reports), and the lack of specific articles in the bibliography (that would discuss the role of specific bacterial species in a condition’s pathogenesis, to support some of the reports’ claims, etc.)

Again, given the various flaws in the data-gathering process, it is assumed that reported findings would not have sufficient scientific validity.

Regarding scientific rigour, some comments from the evaluating model about the reports included the following shortcomings: the lack of appropriate discussion around the study’s limitations, the lack of references about butyryl-CoA CoA-transferase overexpression in *Bacteroides fragilis*, the lack of robust methodologies, the specific mechanisms underlying observed gene dysregulation and metabolic changes were not fully elucidated, the lack of case studies to illustrate discussions, the lack of research on the gene expression profiles of gut bacteria in Rheumatoid Arthritis patients, the fact that some important claims seemed to be unsupported (for example, the increased abundance of *Bacteroides fragilis* in hypertensive individuals and its role in butyrate biosynthesis, which was in turn highlighted as playing a role in blood pressure regulation).

In summary, the reports again scored high for the quality of their writing, but scientific rigour and bibliographic reference quality suffered considerably. Interestingly, one-shot reports were praised for their discussion of study limitations, while stitched reports were found not to contain those discussions. This may be the result of splitting the report-drafting process into six separate components, which meant that the LLM only had a complete view of the text during the last API call, where the context window capacity may have played a role in not allowing for further text-element additions.

## Further Discussion

The results presented here can be said to be the output of a very specific combination of inputs and prompts fed to a specific version of an LLM (itself a black box), which makes it hard to claim that any of the results are definitive or somehow generalizable to the realm of Large Language Models and science. Nonetheless, the final state of our architecture (including prompts and other rules-based algorithms) is the result of a dialogical process that was followed to improve the quality of outputs and to avoid observed vices in the throughput.

While it is difficult to generate evidence that the current form of our LLM prompts and other elements of our architecture is the most suitable for the task at hand, we claim that the current level of quality attained in the script’s throughput and output proved hard to improve further (this is consistent with findings from Liang *et al.* (2023) [18]) or the way to do so was not immediately clear to us. Also, capturing the entire prompt-engineering process as well as the input-output pairs that we produced during the development phase proved a very long task that would have been very complex to order and present in any meaningful way.

However, the script is not only available for reproducibility purposes, but also the initial user parameters and all script-generated throughput for each one of our 5 final runs were saved as Python pickle files ^1^, so that the entire process can be traced from the beginning until the end, and scripts can be re-run from any particular point of the process onwards, both with the current prompts or with improved versions thereof.

An initial approach to generate reports directly from a single all-in-one LLM prompt was abandoned soon due to the impossibility of inspecting the intermediate steps the LLM took inside of the black box and to spot where the process started deviating from an expected path.

Therefore, our approach, while likely imperfect in several ways, it provides transparency regarding inputs and outputs at every single step of the process.

In the below paragraphs, we describe how research could be continued towards our initial objectives.

The fact that we necessarily summarize each article to its most basic components (as in, a table with certain elements) means that important methodological considerations from each study are not passed along to the model for the drafting process. This means that the model drafts the final report starting with a very limited understanding of the research underlying the consolidated data and potentially ignoring important details that would compromise the compatibility of results from across different studies. Also, the models’ limited context window means that the entire body of relevant literature cannot be input while also expecting a very lengthy and deep output.

Thus, we consider it essential to explore whether using a microbiology-trained embeddings model could help improve data extraction from articles, in turn allowing for a process that could feed LLMs with all relevant information in a very concise manner, further allowing them to produce longer, more relevant output.

An alternative to the above would be to explore a way to integrate GPT-4o (or a similar model) with an automated pipeline that would download all articles and relevant data and feed them to the model (without relying on the API’s browsing tool, which the model’s API may have been programmed to decline usage at user’s request). GPT-4o seems to possess more powerful data-extraction capabilities but often declines to review content from provided URLs.

Also, in addressing the model’s shortcomings with scientific rigour, we suggest testing whether instead of focusing on splitting the process into separate report components (Introduction, Discussion, Conclusions, Bibliography, etc), it would produce better results to split the methodology into more granular steps (more related with the scientific process in general, and with the differential abundance analysis process in particular). Steps would have to mimic those followed by researchers when producing similar analyses, and the models’ output at each stage would be equally evaluated for appropriateness.

Further research is also needed on how to engrain in LLMs behaviour the checks and balances that are necessary for scientific rigour. These checks could include the use of references to support claims, exhaustiveness, and transparency about the steps followed to digest data and reach conclusions.

In more general terms, more research is needed regarding the way that LLMs produce output that requires iterative layers or steps of logical reasoning. This would lay the basis for building more accurate prompts and pipelines. It is currently unclear whether an LLM would follow typical steps involved in producing an output. That is, although models can precisely enumerate steps required to produce different types of outputs, it is unclear whether LLMs themselves follow such steps in producing outputs, or what exact logical skills LLMs possess [40]. Understanding such underlying logic and limitations will be key in producing pipelines capable of more complex reasoning.

## Conclusions

In this dissertation, we explored the application of Large Language Models (LLMs) for differential abundance analysis in microbiome research. The rapid advancements in natural language processing (NLP) and artificial intelligence have opened new avenues for automating and enhancing various aspects of scientific research, including the analysis of complex biological data. Our study aimed to assess the capabilities and limitations of LLMs in handling differential abundance tasks, which are crucial for understanding the relationships between microbial communities and various health conditions.

### 6.0.1 Summary of Findings

The research presented in this dissertation highlights several key findings:

1. **Feasibility of LLMs in Differential Abundance Analysis**: The use of LLMs demonstrated potential in automating parts of the differential abundance analysis process. These models can extract relevant information from scientific literature, generate summaries, and suggest hypotheses based on existing data. However, different models show considerably different levels of skill, with GPT-4o being the most accurate for this matter, although hard to integrate to an automated pipeline, due to GPT-4o’s API proclivity to decline to review external internet sites.
2. **ChatGPT and Gemini not a reliable source of academic literature**: Webscraping extraction of sources was successful and could be a good first step in any additional efforts to reproduce a similar pipeline. However, many research journals are not robot-friendly, thus reducing the number of useful sources of data.
3. **Varying quality and availability of data limits our pursuit**: Many relevant articles do not contain supplementary data or available data is not complete, is hard to extract, or does not contain sufficient metadata to reproduce results.
4. **Challenges with Data Accuracy and Consistency**: Despite their potential, LLMs faced significant challenges in producing accurate and consistent results. Issues such as hallucinations, incomplete data extraction, and the generation of incorrect references were common. These challenges underscore the need for careful validation and cross-checking of LLM-generated outputs in scientific research. It is important to include evaluation steps to verify output and throughput along the generative process.
5. **Impact on Scientific Rigor**: The study found that while LLMs can assist in drafting reports and summarizing data, they often fall short in maintaining the scientific rigor required for high-quality research. The generation of references, the synthesis of complex data and the ’thought process’ behind the conclusion-extraction process in LLMs were particularly problematic, leading to concerns about the reliability of LLM-based research outputs.
6. **Context windows may be still insufficient**: While context windows have been growing importantly in the recent past, their current proportions still limit the amount of data that can be fed into models and the amount (and thus depth) of texts that are generated from it.
7. **LLMs may need less freedom**: The researcher’s role in chunking complex research processes into irreducibly accurate steps with outputs for which LLMs have a known and positive track record, may prove essential.
8. **Potential for Integration with Human Expertise**: One of the key takeaways from this research is the potential for integrating LLMs with human expertise. While LLMs can handle repetitive and time-consuming tasks, human oversight remains essential to ensure the accuracy and scientific validity of the results.
9. **Skill differentiation in LLMs**: It is important for researchers and users to understand the difference between a skill to enumerate steps needed to produce an output, and the skills needed to produce that same output. With LLMs’ ability to recite detailed lists of almost anything, it is tempting to think they are ready to produce any type of output.

### 6.0.2 Implications for Future Research

The findings from this dissertation suggest several directions for future research:

1. **Improving LLM Accuracy and Consistency**: There is a need for continued development of LLMs to improve their accuracy and consistency, particularly in the context of scientific research. This includes refining algorithms to reduce hallucinations and improve the quality of data extraction and synthesis.
2. **Hybrid Models Combining AI and Human Expertise**: Future research could explore the development of hybrid models that combine the strengths of LLMs with human expertise. Such models could leverage the efficiency of AI while ensuring the scientific rigor of human-driven research.
3. **Expanding the Scope of LLM Applications**: While this study focused on differential abundance analysis, LLMs have the potential to impact various other areas of microbiome research and beyond. Future studies could explore their application in other complex biological data analyses.

### 6.0.3 Final Thoughts

The integration of LLMs into scientific research represents a significant advancement in the field of computational biology. However, as this dissertation has shown, there are still considerable challenges to overcome before LLMs can be fully trusted to perform complex scientific tasks independently. The findings of this research contribute to the ongoing conversation about the role of AI in science and highlight the importance of continued innovation and evaluation considerations in the development and application of these technologies.

In conclusion, while LLMs offer exciting possibilities for enhancing research efficiency and productivity, their limitations necessitate a cautious approach. The future of AI-assisted research lies in the collaboration between human researchers and AI, where each complements the other’s strengths to achieve reliable, rigorous, and impactful scientific outcomes.

## Footnotes

1 https://gemini.google.com/

2 https://chat.openai.com/

3 https://www.mg-rast.org/

4 https://www.ebi.ac.uk/metagenomics

5 data could be found in a number of formats including tables in PDF file, word documents, graphs, Excel spreadsheets or links to external websites that required additional steps to find.

6 term used to describe claims often made by LLMs which often sound realistic but are misleading or outright made up.

7 https://www.biorxiv.org/

8 the ”short term” memory of the model.

1 Refer to Appendix A.2 for details on how to do this

1 Refer to Appendix A.2

1 https://chatgpt.com/g/g-HMNcP6w7d-data-analyst

