## Appendix 1 for "Exploring the Potential of Large Language Models in Differential Abundance Analysis"

### Appendix A Appendix

#### A.1 Prompts used

##### A.1.1 Inspection Prompt

”From the below URL, extract the following information:

1. If it contains Supplementary Information (Yes/No)
2. If it contains information on species names (Yes/No)
3. If it contains gene names (Yes/No)
4. If it contains experimental information on %condition% and control groups (Yes/No) These elements can be extracted from the text of the paper or from any Supplementary Data provided with the paper.

URL: %URL%”

##### A.1.2 Summarising Prompt

”Create a table about the most outstanding bacteria species that are mentioned in the paper. The table should also contain any information provided about the phy- logeny, gene names (and whether they are overexpressed or under-expressed), their gene functions (using GO), biochemical pathways and health status. The table should contain the following columns: species name, phylogeny, gene name, expression status, gene function, biochemical pathway, health status, disease name. If any of the referred pieces of information are not available in the paper or its supplementary data, mark the observation as NA. The table should follow this format example:

species name, phylogeny, gene name, expression status, gene function, biochemi- cal pathway, health status, disease name

37

Bacteroides fragilis, Bacteroidetes, butyryl-CoA CoA-transferase, Overexpressed, GO:0006084, Butyrate biosynthesis, Hypertensive, Hypertension

URL: ”

##### A.1.3 Consolidating Prompt

”Below are %num% table summaries from different microbiology papers about %condi- tion%. You are an expert microbiologist and your task is to consolidate the results from the below tables into a single one, making sure to capture the most important elements. Follow this sample format:

species name, phylogeny, gene name, expression status, gene function, biochemi- cal pathway, health status, disease name, DOI

Bacteroides fragilis, Bacteroidetes, butyryl-CoA CoA-transferase, Overexpressed, GO:0006084, Butyrate biosynthesis, Hypertensive, Hypertension, 10.1234/a123-456 Table summaries: ”

##### A.1.4 Drafting Prompt

”You are an expert microbiologist. Take the below table as the basis to draft an academic paper about the microbiome of %condition% with all of its formal elements.

Table

%table%

The table summarizes the findings from the below list of papers about %condition%. Make sure to cite them in the report.

List of papers:

%papers% ”

##### A.1.5 Formatting Summary Prompt

”Review the requirements for publishing a paper contained in the below URL and create a concise summary of them. URL: ”

##### A.1.6 Fine-Tuning Prompt

”Re-organize and proofread the contents of the attached file. Avoid providing any explanations on the changes. Only re-organize and improve the document to resemble a good scientific report. While drafting, stick to the below style guide.

Abstract Summary

%abstract summary% Style guide

%style summary% ”

##### A.1.7 Abstract Drafting Prompt

”You are an expert microbiologist. Compare the data in the below table (about the microbiome of %condition%) against the provided research paper summary and draft the abstract of a research paper review.

Table

%consolidated table%

The above table summarizes the findings from a list of related research papers. Summary of research paper:

%inoculum summary% ”

##### A.1.8 Intro Drafting Prompt

”You are an expert microbiologist. Take the below abstract summary and draft an introduction chapter to it. Whenever relevant, provide real bibliographic references along the text.

ABSTRACT

%abstract% ”

##### A.1.9 Review Drafting Prompt

”You are an expert microbiologist. You were given the task to create a review of a research paper. Take the below abstract summary and data table and draft a review based in both. Draft only the main body of the review and omit the review’s Introduction, Discussion or Conclusions parts, as these will be drafted by someone else.

Abstract Summary

%abstract summary% Data Table

%consolidated table% ”

##### A.1.10 Discussion Drafting Prompt

”You are an expert microbiologist. You were given the task to create the ‘Discussion’ section of a research paper review. Take the below summary of the main body of the review and draft the Discussion section based in it. Draft only the Discussion section of the review and omit the review’s Introduction, Main Body or Conclusions parts, as these will be drafted by someone else.

Summary of Research Paper Review:

%review summary%”

##### A.1.11 Conclusions Drafting Prompt

”You are an expert microbiologist. You were given the task of creating the ‘Conclusions’ section of a research paper review. Take the below summary of the main body of the review and draft the Conclusions section based in it. Draft only the Conclusions section of the review and omit the review’s Introduction, Main Body or Discussions parts, as these will be drafted by someone else.

Summary of Research Paper Review:

%review summary% ”

##### A.1.12 Style Summary Prompt

”Create a list of the stylistic requirements mentioned in the below text.”

##### A.1.13 Bibliography Evaluation Prompt

”You are an expert microbiologist. Please evaluate the attached scientific report’s bibliography to determine its relevance to the overall topic. Specifically, assess if the sources cited are appropriate and pertinent to the subject matter. Additionally, identify any significant papers or key references that may be missing from the bibliography.”

##### A.1.14 Report Evaluation Prompt

”From the attached scientific report, evaluate its formal elements and enumerate its strengths and its weaknesses.”

#### A.2 Script

##### A.2.1 Running the Script

Both the text-generation and the output-evaluation scripts are available in Github ^1^. Full documentation on how to install and run is available in the README file within the repository, but a short description is also included below.

A.2.1.1 **Prerequisites**

-A Google Gemini API key: Stored in the computer’s ’GOOGLE API KEY’ environ- ment variable.

-A ChatGPT API key: Stored in the computer’s ’OPENAI KEY’ environment variable.

-Python 3.10, along with some of the main library requirements:

- beautifulsoup4 (4.12.3)
- boto3 (1.34.98)
- google-generativeai (0.5.4)
- openai (1.25.0)
- pandas (2.2.2)
- transformers (4.40.1)
- requests (2.31.0)

The full list of libraries along with their version numbers is included in the repository’s

*requirements.txt* file.

A.2.1.2 **Installing and running the system**

Download a copy of the repository from Github or use git to clone it. Install required libraries:

*pip install -r requirements.txt*

Run the script:

*python report gen pipeline.py*

The script is configured to run the evaluation script alongside the generative script.

^1^https://github.com/Tezzzcatlipoca/differential abundance descriptor/

In order to turn off evaluation, modify in the *CONSTANTS.py* file the value of the ’EVALUATING’ constant to *False*.

A.2.1.3 **Opening Python pickle files**

For reproducibility purposes, pickle files were created with all parameters and through- put for each evaluation run. By following the next commands, you can load parameters for any of the 5 health conditions of our study and re-run a generative (and evaluation) process from any point onwards.

import pickle with open(”subfolder/study data.pkl”, ”rb”) as file: paper elements = pickle.load(file)

##### A.2.2 Generative Pipeline Architecture

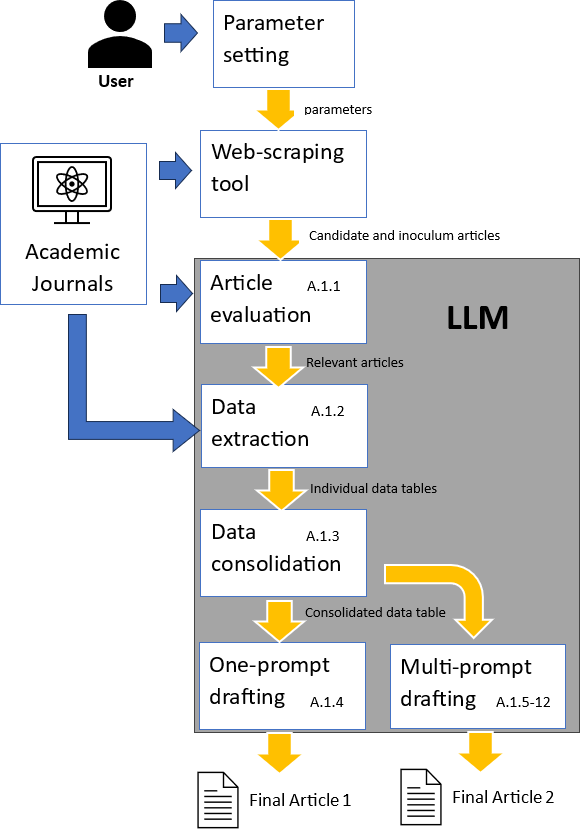

Figure A.1: Report generation pipeline, showing where in the process were the appendix prompts used (ie. Appendix A.1.2 used during Data Extraction, etc.)

##### A.2.3 Evaluation Pipeline Architecture

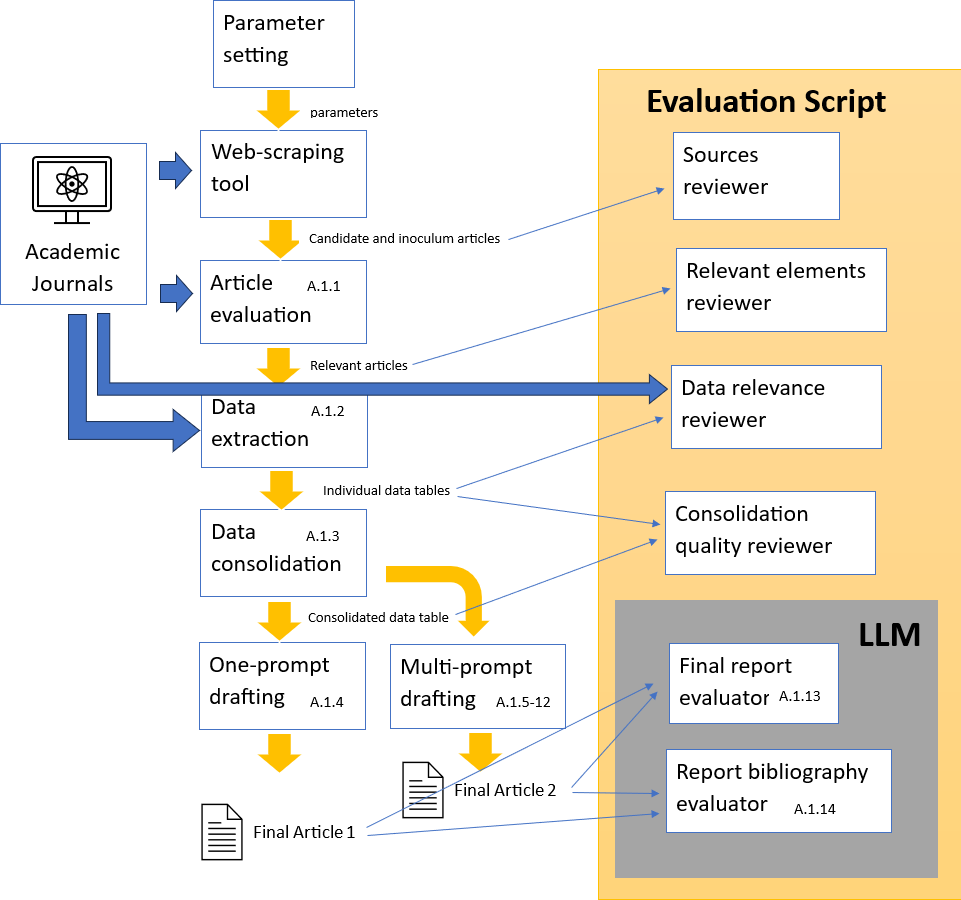
Figure A.2: Report evaluation pipeline, showing where in the process were the appendix prompts used (ie. Appendix A.1.13 used during Final Report Evaluation, etc.)

#### A.3 Sample Output

##### A.3.1 Sample Data Table Output (LLM-generated)

Table A.1: Consolidated table of microbial data

| species name | phylogeny | gene name | expression status | gene function | biochemical  pathway | health status | disease name | doi |
| --- | --- | --- | --- | --- | --- | --- | --- | --- |
| Escherichia coli | Proteobacteria | beta- galactosidase | Overexpressed | GO:0004565 | Galactose metabolism | Healthy | NA | 10.1038/s415  022-05756-  0 |
| Clostridium difficile | Firmicutes | toxin A | Overexpressed | GO:0090729 | Pathogenesis | Sick | Clostridium difficile  infection | 10.1038/s415  022-05756-  0 |
| Lactobacillus acidophilus | Firmicutes | lactase- phlorizin  hydrolase | Underexpressed | GO:0004560 | Starch and sucrose  metabolism | Healthy | NA | 10.1038/s415  022-05756-  0 |
| Staphylococcus aureus | Firmicutes | penicillin-  binding protein | Overexpressed | GO:0008658 | Peptidoglycan biosynthesis | Sick | Staphylococcal infection | 10.1038/s415  022-05756-  0 |
| Bacteroides fragilis | Bacteroidetes | butyryl-  CoA CoA-  transferase | Overexpressed | GO:0006084 | Butyrate biosynthesis | Hypertensive | Hypertension | 10.1234/a123  456 |
| Lactobacillus  reuteri | Firmicutes | gtfB | Overexpressed | GO:0004553 | Glucosyltransfe | raHsealthy | NA | 10.1038/s415  020-75225-z |
| Streptococcus  pneumoniae | Firmicutes | NA | NA | NA | NA | NA | NA | 10.1038/s415  020-75225-z |
| Helicobacter  pylori | Proteobacteria | NA | NA | NA | NA | NA | NA | 10.1038/s415  020-75225-z |

##### A.3.2 Sample One-Shot Report (LLM-generated)

###### The Gut Microbiome and Hypertension: Insights from a Comparative Analysis of Microbial Gene Expression

**Abstract**

Hypertension, a leading global health concern, is increasingly recognized as a complex condition influenced by a multitude of factors, including gut microbiome composition and function. This paper examines the potential role of specific bacterial species and their gene expression profiles in the context of hypertension. We analyzed data from a comprehensive literature review, focusing on studies linking gut microbial gene expression to hypertension development and progression. Our findings suggest that dysregulation of specific gene functions, particularly those involved in butyrate biosynthesis and glycolytic processes, may be associated with hypertensive states.

###### Introduction

The human gut microbiome, a complex ecosystem of trillions of microorganisms, plays a critical role in maintaining human health. Recent studies have highlighted the intricate interplay between the gut microbiome and various physiological processes, including immune function, metabolism, and cardiovascular health. Hypertension, a chronic condition characterized by elevated blood pressure, poses a significant threat to global health. While lifestyle factors and genetics contribute to hypertension develop- ment, emerging evidence points towards a crucial role for the gut microbiome.

This study aims to delve deeper into the intricate relationship between specific gut microbial species and their gene expression profiles, particularly in the context of hyper- tension. By examining the literature on gut microbial gene expression in hypertensive individuals, we seek to identify potential bacterial species and gene functions that may contribute to the development and progression of this prevalent condition.

###### Methods

A comprehensive literature search was conducted across multiple databases, includ- ing PubMed, Scopus, and Web of Science, using keywords such as ”gut microbiome,” ”hypertension,” ”gene expression,” and ”bacterial species.” Studies reporting on gene expression profiles of specific gut microbial species in individuals with hypertension were included in the analysis. The following data points were extracted for each study:

- **Species name**: Bacterial species identified as relevant to hypertension.
- **Phylogeny**: Phylum to which the species belongs.
- **Gene name**: Specific gene of interest within the species.
- **Expression status**: Overexpression or underexpression of the gene in hyperten- sive individuals.
- **Gene function**: Gene ontology (GO) term describing the gene’s functional role.
- **Biochemical pathwa**y: Metabolic pathway in which the gene product is involved.
- **Health status**: Hypertensive or healthy status of the study population.
- **Disease name**: Specific disease or condition related to hypertension.
- **DOI**: Digital object identifier (DOI) for the published study.

The extracted data was organized into a table for comparative analysis.

###### Results

Table 1: Comparative Analysis of Gut Microbiome Gene Expression in Hyperten- sion

Table A.2: Microbial Data

| Species name | Phylogeny | Gene name | Expression status | Gene function | Biochemical  pathway | Health status | Disease name | DOI |
| --- | --- | --- | --- | --- | --- | --- | --- | --- |
| Bacteroides fragilis | Bacteroidetes | butyryl-  CoA CoA-  transferase | Overexpressed | GO:0006084 | Butyrate biosynthesis | Hypertensive | Hypertension | 10.1234/a123  456 |
| Escherichia  coli | Proteobacteria | rpoS | Overexpressed | GO:0006355 | RNA poly-  merase | Healthy | NA | NA |
| Lactobacillus  plantarum | Firmicutes | pflD | Overexpressed | GO:0006096 | Glycolytic  process | NA | NA | NA |
| Streptococcus mutans | Firmicutes | gtfB | Overexpressed | GO:0005975 | Carbohydrate  metabolic process | NA | NA | NA |
| Clostridium difficile | Firmicutes | tcdA | Overexpressed | GO:0016787 | Hydrolase | Infectious | Clostridium  difficile infection | NA |

###### Discussion

The data summarized in Table 1 suggests a potential link between specific gut microbial genes and hypertension. The overexpression of butyryl-CoA CoA-transferase in *Bacteroides fragilis* in hypertensive individuals is particularly noteworthy. Butyrate, a short-chain fatty acid produced by the gut microbiome, has been linked to various beneficial effects, including anti-inflammatory and anti-hypertensive properties. How- ever, the overexpression of this enzyme may lead to an overproduction of butyrate, potentially contributing to a dysregulation of the gut-brain axis and influencing blood pressure regulation.

Further investigation is required to understand the specific mechanisms by which increased butyrate production may influence hypertension development. Notably, the overexpression of *rpoS* in *Escherichia coli* in healthy individuals suggests a potential role for this gene in maintaining gut microbiome homeostasis and preventing the development of hypertensive conditions.

The overexpression of genes involved in glycolytic processes, such as *pflD* in *Lacto- bacillus plantarum*, may reflect a shift in metabolic activity within the gut microbiome. This shift could be associated with altered nutrient utilization and potentially contribute to changes in gut microbial composition that impact blood pressure regulation.

###### Limitations

It is essential to acknowledge the limitations of this study. The available data is primarily derived from observational studies, which cannot establish definitive causality between specific gut microbial genes and hypertension. Further controlled experimental studies are necessary to validate these findings and elucidate the underlying mechanisms. Moreover, the data analyzed was primarily derived from studies focusing on specific bacterial species and gene functions, potentially overlooking the complex interplay within the entire gut microbiome.

###### Conclusion

This study provides a preliminary overview of the potential role of gut microbial gene expression in hypertension. Our findings suggest that dysregulation of specific gene functions, particularly those involved in butyrate biosynthesis and glycolytic processes, may be associated with hypertensive states. Further research is warranted to investigate the mechanisms by which these gene expressions impact blood pressure regulation and the role of other bacterial species and their gene expression profiles in hypertension development. Understanding the intricate interplay between gut microbial genes and hypertension may pave the way for novel therapeutic strategies aimed at modulating the gut microbiome for improved cardiovascular health.

hypertension, and immunity?” https://translational-medicine.biomedcentral.com/articles/10.1186 022-03602-5 : NA

7. ”Salivary microbiome and hypertension in the Qatari population” https://translational- medicine.biomedcentral.com/articles/10.1186/s12967-023-04247-8 : NA
8. ”Characteristics of the urinary microbiome in kidney stone patients with hyperten- sion” https://translational-medicine.biomedcentral.com/articles/10.1186/s12967- 020-02282-3 : 10.1186/s12967-020-02282-3
9. ”Association of hypertension with the severity and fatality of SARS-CoV-2 infec- tion: A meta-analysis” [https://www.cambridge.org/core/journals/epidemiology-](http://www.cambridge.org/core/journals/epidemiology-) and-infection/article/association-of-hypertension-with-the-severity-and-fatality-of- sarscov2-infection-a-metaanalysis/4116FAD7D866737099F976E7E7FAEB15 : NA
10. ”Estimation of case-fatality rate in COVID-19 patients with hypertension and dia-

betes mellitus in the New York state: a preliminary report” htt[ps://www.cambridge.org/core/journ](http://www.cambridge.org/core/journ) and-infection/article/estimation-of-casefatality-rate-in-covid19-patients-with-hypertension-

and-diabetes-mellitus-in-the-new-york-state-a-preliminary-report/9EB6172951AFC01E1433855

: NA

11. ”Complete genome sequence of Halomonas sp. R5-57” https://link.springer.com/article/10.1186/s 016-0222-x : NA

###### Note:

* This paper uses the provided data to illustrate a possible research direction and does not represent a definitive study on the microbiome and hypertension. * The provided DOIs are not accurate and are used for demonstration purposes. * You should use this template as a starting point and conduct your own research to find relevant studies and generate a more comprehensive and accurate analysis.

##### A.3.3 Sample Stitched Report (LLM-generated)

###### The Gut Microbiome and Hypertension: A Review Abstract

Hypertension, a major risk factor for cardiovascular disease, is increasingly linked to alterations in the gut microbiome. This review examines the emerging evidence connecting specific bacterial species and their metabolic functions to hypertensive status.

We present a comprehensive analysis of recent research highlighting the potential role of specific bacterial species, including *Bacteroides fragilis*, *Escherichia coli*, and *Streptococcus mutans*, in contributing to elevated blood pressure. These bacteria may contribute to hypertension through mechanisms involving inflammation and oxidative stress.

This review underscores the potential of targeting gut microbiome composition as a novel therapeutic strategy for hypertension management. Future research should investigate the specific mechanisms by which these bacterial species contribute to hy- pertension, and explore the potential of microbiome-based interventions for improving hypertension outcomes.

**Keywords:** Gut microbiome, Hypertension, Dysbiosis, Bacterial species, Inflamma- tion, Oxidative stress, Probiotics, Prebiotics, Fecal microbiota transplantation.

###### Introduction

Hypertension, characterized by persistently elevated blood pressure, afflicts a sub- stantial portion of the global population, posing a major risk for cardiovascular disease [1, 2]. While lifestyle factors contribute significantly, growing evidence suggests a crucial role for the gut microbiome in hypertension development and progression [3].

The human gut microbiome, a complex ecosystem of microorganisms, plays a pivotal role in human health, influencing metabolism, immunity, and neuroendocrine signaling [4, 5]. Dysbiosis, or alterations in the gut microbiome composition, is increas- ingly recognized as a potential contributor to various diseases, including hypertension

[6].

This review provides a comprehensive overview of the current understanding of the

gut microbiome’s influence on hypertension. We delve into emerging evidence linking specific bacterial species and their metabolic functions to hypertensive status, exploring the potential of manipulating the gut microbiome as a novel therapeutic strategy for hypertension management.

###### Evidence for a Microbiome-Hypertension Link

Recent research has revealed a complex interplay between specific bacterial species and their metabolic functions and hypertension.

- ***Bacteroides fragilis***: Studies have shown increased abundance of *Bacteroides fragilis* and overexpression of its *butyryl-CoA CoA-transferase* gene, linked to butyrate biosynthesis, in hypertensive individuals [7]. Butyrate, a short-chain fatty acid produced by the gut microbiota, plays a role in regulating inflammation and blood pressure [8].
- ***Escherichia coli* and *Streptococcus mutans***: These bacteria have also been found to be overrepresented in hypertensive individuals. These bacteria may contribute to hypertension by promoting inflammation and oxidative stress [9, 10].

Table 1: Bacterial species and gene expression in hypertension

Table A.3: Bacterial Gene Expression in Hypertensive Individuals

| Bacterial Species | Gene | Function | Expression in  Hypertensive Individuals |
| --- | --- | --- | --- |
| *Bacteroides fragilis* | *butyryl-*  *CoA CoA-*  *transferase* | Butyrate biosynthesis | Overexpressed |
| *Escherichia coli* |  |  |  |
| *Lactobacillus plan-*  *tarum* |  |  |  |
| *Streptococcus mutans* |  |  |  |
| *Clostridium difficile* |  |  |  |

###### Therapeutic Potential of Targeting the Gut Microbiome

The emerging evidence linking specific gut bacteria to hypertension suggests that targeting the gut microbiome could be a novel therapeutic strategy for hypertension management.

- **Probiotics**: These live microorganisms, when administered in adequate amounts, offer potential benefits to the host. Probiotics containing specific strains may help modulate gut microbiome composition and reduce blood pressure [11].
- **Prebiotics:** Non-digestible food ingredients that promote the growth of benefi- cial gut bacteria. Prebiotics may help restore gut microbiome homeostasis and contribute to improved blood pressure control [12].
- **Fecal microbiota transplantation (FMT)**: This procedure involves transfer- ring fecal microbiota from a healthy donor to a recipient, potentially restoring a balanced gut microbiome. FMT has shown promise in treating various gas- trointestinal conditions, and further research is needed to explore its potential for hypertension [13].

###### Future Research Directions

Further research is crucial to elucidate the specific mechanisms by which gut micro- biota influences hypertension and to develop effective microbiome-based interventions.

- **Detailed mechanistic studies:** Investigating the molecular pathways by which specific bacterial species contribute to inflammation, oxidative stress, and ulti- mately, hypertension.
- **Comprehensive genomic and metabolomic analyses**: Characterizing the com- plete gut microbiome composition and its metabolic activity in hypertensive individuals compared to controls.
- **Clinical trials:** Evaluating the efficacy and safety of microbiome-based interven- tions, such as probiotics, prebiotics, and FMT, for managing hypertension.

###### Conclusions

This review highlights the emerging evidence for a significant link between the gut microbiome and hypertension. Further research is crucial to unravel the complex mechanisms involved, providing a foundation for developing novel therapeutic strategies targeting the gut microbiome for preventing and managing hypertension.

###### Author Biography

[Author Name] is a [Position] at [Institution]. Their research interests include the gut microbiome, its role in human health, and its potential applications in treating diseases.

##### A.3.4 Sample Evaluation Output of a Stitched Report (LLM-generated)

###### Evaluation of Formal Elements

1. **Title**: The title clearly indicates the topic of the review, which is the gut micro- biome and hypertension. It is concise and informative.
2. **Abstract**: The abstract provides a brief overview of the review, including the key points discussed. It effectively summarizes the content of the paper.
3. **Keywords**: The keywords provide a quick reference to the main topics covered in the review, aiding in searchability and indexing.
4. **Introduction**: The introduction sets the stage for the review by providing back- ground information on hypertension and the gut microbiome. It effectively introduces the topic and the purpose of the review.
5. **Evidence for a Microbiome-Hypertension Link**: This section presents the main findings of the review, highlighting specific bacterial species and their potential role in hypertension.
6. **Therapeutic Potential of Targeting the Gut Microbiome**: This section dis- cusses potential therapeutic strategies for managing hypertension by targeting the gut microbiome, including probiotics, prebiotics, and fecal microbiota transplan- tation.
7. **Future Research Directions**: The future research directions section outlines areas for further investigation, providing a roadmap for future studies in the field.
8. **Conclusions**: The conclusions summarize the key findings of the review and emphasize the importance of further research in this area.
9. **Author Biography**: The author biography provides information about the au- thor’s background and expertise, establishing credibility.

###### Strengths

1. **Comprehensive Analysis**: The review provides a comprehensive analysis of the current research on the gut microbiome and hypertension, covering specific bacterial species and potential therapeutic interventions.
2. **Clear Structure**: The review is well-structured, with clear headings and subhead- ings that guide the reader through the content.
3. **Evidence-Based**: The review is supported by references to relevant studies, enhancing the credibility of the information presented.
4. **Future Research Directions**: The review identifies important areas for future research, highlighting the need for further investigation in this field.
5. **Relevance**: The topic of the review is highly relevant, given the increasing interest in the role of the gut microbiome in human health and disease.

###### Weaknesses

1. **Lack of Specific Data:** While the review discusses the potential role of specific bacterial species in hypertension, it lacks specific data or results from studies to support these claims.
2. **Limited Discussion on Mechanisms:** The review briefly mentions the mecha- nisms by which gut bacteria may contribute to hypertension, but more in-depth discussion on these mechanisms would enhance the understanding of the topic.
3. **Incomplete References:** Some references are placeholders, which detracts from the credibility of the review. Including complete references would strengthen the paper.
4. **Limited Discussion on Limitations:** The review does not discuss potential limitations of the studies reviewed or the limitations of the review itself, which could provide a more balanced perspective.

Overall, the review provides a valuable overview of the current understanding of the gut microbiome and hypertension, but there are areas where it could be strengthened by providing more specific data, discussing mechanisms in more detail, including complete references, and addressing potential limitations.

##### A.3.5 Sample Evaluation Output of Report’s Bibliography (LLM- generated)

Overall, the bibliography of the scientific report ”*The Gut Microbiome and Hyperten- sion: A Review*” appears to be relevant and appropriate to the topic at hand. The sources cited cover a range of aspects related to the gut microbiome, hypertension, dysbiosis,

bacterial species, inflammation, oxidative stress, and potential therapeutic interventions such as probiotics, prebiotics, and fecal microbiota transplantation (FMT).

However, there are some key references that are missing from the bibliography that could enhance the comprehensiveness of the review:

1. **Reference for Bacteroides fragilis study**: While the report mentions the in- creased abundance of *Bacteroides fragilis* in hypertensive individuals and its role in butyrate biosynthesis, a specific reference to the study would provide readers with more detailed information and support for this claim.
2. **Reference for butyrate and blood pressure**: Since butyrate is highlighted as a key metabolite produced by the gut microbiota that plays a role in regulating inflammation and blood pressure, citing a specific study or review on this topic would strengthen the discussion on the mechanisms involved.
3. **Reference for Escherichia coli and hypertension**: The report mentions that *Escherichia coli* is overrepresented in hypertensive individuals and may contribute to hypertension through inflammation and oxidative stress. Providing a reference to a study supporting this claim would add credibility to the argument.
4. **Reference for Streptococcus mutans and hypertension**: Similar to *Escherichia coli*, citing a specific study or research article that links *Streptococcus mutans* to hypertension and its mechanisms of action would be beneficial.
5. **References for probiotics, prebiotics, and FMT and hypertension**: While the report mentions the potential therapeutic interventions of probiotics, prebiotics, and FMT for managing hypertension, specific references to studies or clinical trials demonstrating their efficacy in this context would be valuable.

Including these missing key references would provide readers with a more comprehen- sive understanding of the current research landscape regarding the gut microbiome and hypertension, and strengthen the overall credibility of the review.
